# Quantitative polymerase chain reaction from malaria rapid diagnostic tests to detect *Borrelia crocidurae* in febrile patients in Mali

**DOI:** 10.64898/2026.02.13.705696

**Authors:** Pascal Dembélé, Adama Zan Diarra, Papa Mouhamadou Gaye, Agokeng Dongmo Armel Joseph, Li Bing, Mahamadou Ali Thera, Stéphane Ranque

## Abstract

Tick-borne relapsing fever (TBRF) is caused by *Borrelia* species transmitted to humans by soft ticks of the genus *Ornithodoros*. Very little is currently known about the morbidity of this disease in Mali, despite the risk of human co-infection with malaria. The lack of appropriate diagnostic services or technical expertise to differentiate suspected malaria from other causes of febrile illness of unknown origin means that this tick-borne disease remains neglected and under-diagnosed in febrile patients in Mali. Our study investigated the detection of *Borrelia crocidurae* DNA in febrile patients in Mali from malaria rapid diagnostic tests (RDTs).

**Methods:** Between June and December 2021, negative and positive malaria RDTs were collected from 41 sites in the nine regions of Mali. Both qPCR and standard PCR were used to detect the presence of *B. crocidurae* DNA.

**Results:** Of the 1496 malaria RDTs tested, *B. crocidurae* DNA was detected in 9 malaria-negative RDTs. All of these were collected in the region of Kayes, where the prevalence rate was 6% (9/150) of the negative malaria RDTs.

**Conclusions:** This study demonstrates that tick-borne relapsed fever is an under-diagnosed condition in febrile patients in Mali.

## Introduction

Transmitted recurrent fevers (TBRF) are neglected febrile infections caused by soft ticks of the genus *Ornithodoros* spp .[1, 2]. Wild rodents and insectivores are common reservoir hosts. They are caused by several spirochetes of the genus *Borrelia* (*B. crocidurae, B. duttonii, B. recurrentis, and B. hispanica*), which are endemic in subtropical regions worldwide [3–5]. This acute febrile illness causes multiple recurrences of nonspecific signs and symptoms, including fever, headache, hepatomegaly, splenomegaly, anemia, myalgias, and arthralgias. These symptoms are similar to those of malaria [6,7]. The first clinical cases of (TBRF) have long been described as a major cause of morbidity and mortality in many parts of Africa [8,9]. In West Africa, *B. crocidurae* is recognized as the only vector responsible for (TBRF) disease, especially in many rural populations living in extremely precarious conditions [10,11]. Studies have shown that the incidence of TBRF in West Africa is high, accounting for 13% of febrile illnesses treated in rural health facilities [2]. In Tanzania, particular attention is currently being paid to TBRF, which is one of the ten leading causes of death in children under 5 year of age [12].

Microscopy is used to diagnose the disease. It is usually based on observation of the spirochete in thick blood smears during the acute febrile phase, a technique identical to that used to detect malaria haematozoa [13]. On the other hand, molecular methods (PCR or qPCR) remain the best diagnostic tools because of their high sensitivity and specificity [14,15]. The lack of diagnostic tools in Africa has been identified as a major gap in healthcare delivery on the continent [16]. In Mali, for example, studies have reported the presence of *B. crocidurae* in tick vectors and small mammal rodents, which are the reservoirs [17–19]. It has been shown that West African tick-borne relapsing fever (TBRF) is widespread, especially in the Saharan, Sahelian and Sudano-Sahelian regions [3], where average annual precipitation is between 50 and 250 mm, 250 and 500 mm and 500 and 750 mm respectively [3]. The persistence of drought in sub-Saharan countries potentially contributes to the proliferation of *Ornithodoros sonrai* responsible for *B. crocidurae* [20,21]. Most studies on TBRF in West Africa have been carried out in Senegal [14,22,23], where the TBRF vector ticks are geographically distributed over the northern two-thirds of the country, north of the 750 mm isohyet, and the southern limit of the vector tick corresponds approximately to latitude 13°40’N [3].

However, TBRF remains the second challenge for many malaria-endemic African countries, including Mali, where blood smears are tested only for Plasmodium in blood [24]. Because of the rarity of appropriate diagnostic services, such as molecular biology laboratories, it is clear that TBRF cannot be diagnosed quickly and reliably [23]. Indeed, when misdiagnosed, many cases of TBRF will be considered treatment-resistant malaria, as studies in Togo have shown, where febrile patients were often misdiagnosed as malaria [6]. Rapid diagnostic tests (RDTs) remain the most used tool in healthcare establishments, due to their ease of use. However, several studies have pointed to their usefulness as a potential source of DNA for molecular analysis[7,25–27]. Therefore, we used DNA extracted from RDTs to determine the prevalence of TBRF as a cause of fever in RDT-negative patients in Mali.

## Materials and methods

### Study sites

Mali is a landlocked country in West Africa located between the 10- and 25-degree north latitudes and between the 4- and 12-degree west longitudes. It covers a 1,241,238 km2 area. There are 3 climatic zones in Mali that extend from south to north: the Sudano-Guinean zone, which covers 25% of the territory and has a rainfall of approximately 1300 to 1500 mm per year; the Sahelian zone, which covers 50% of the territory and receives rainfall of 200 to 800 mm per year; and the Saharan desert zone, which represents 25% of the territory. This zone is marked by irregular rainfall, often less than 200 mm per year. For this study, we collected RDT cassettes from 41 sites in the following regions of Mali: Kayes, Koulikoro, Sikasso, Ségou, Mopti, Timbuktu, Gao, Kidal and Menaka, from June to December 2021. No sample was collected in the Taoudeni region or in the capital city Bamako.

### Borrelia spp. real-time PCR

DNA extraction for *Borrelia* detection from malaria RDTs was performed as previously described [29]. The DNA templates were subjected to real-time polymerase chain reaction (qPCR) using primers and probes targeting the internal transcribed spacer (ITS) region of the *Borrelia* spp. rRNA gene. The primer sequences used were Bor_ITS4_F 5’-GGCTTCGGGTCTACCACATCTA-3’ and Bor_ITS4_R 5’-CCGGGAGGGGAGTGAAATAG-3’; and the Bor_ITS4_ P probe sequence was 6FAM-TGCAAAAGGCACGCCATCACC TAMRA [18]. The samples that tested positive for *Borrelia* spp. by ITS PCR were subjected to a second qPCR targeting the glpQ gene specific for B. crocidurae using the primers (croci_glpQ_F5’-CCTTGGATACCCCAAATCATC-3’ and croci_glpQ_R5’-GGCAATGCATCAATTCTAAAC-3’) and the B. croci_glpQ_P probe (6FAM-ATGGACAAATGACAGGTCTTAC -MGB) [30, 31]. The qPCR mix composition and reaction steps were the same as those previously described by Keita et al [33]. To validate our results, each PCR analysis was performed with negative controls and a positive control derived from DNA of *B. crocidurae* strains grown in the laboratory. Samples were considered positive if the q-PCR cycle threshold (Ct) was less than 36 [32].

### DNA sequencing

All samples positive for the two genes by qPCR were subjected to standard PCR and sequencing. Standard PCR amplification targeting the Flagellin *flaB* gene (B_F1 (5’-TAATACGTCAGCCATAAATGC-3’ and B_R1 (5’-GCTCTTTGATCAGTTATCATTC-3’) [14] was performed using a thermocycler (Applied Biosystems, Foster City, CA, USA). The composition of the mixture and the standard PCR program and sequence were the same as those previously described by Rahal M. et al [34]. The amplified products were purified using a Macherey Nagel plate (NucleoFast 96 PCR, Düren, Germany) and sequenced using the same primers (B_F1 and B_R1). Sequencing was performed using the same primers as for PCR with BigDye Terminator v1.1, v3.15× sequencing buffer (Applied Biosystems, Warrington, UK) and performed on an ABI 3100 automated sequencer (Applied Biosystems). The resulting sequences were assembled and processed using ChromasPro (www.technelysium.com.au/chromas.html) [35] and aligned using BioEdit software (http://www.mbio.ncsu.edu/BioEdit/bioedit.html). The corrected sequences were compared with existing sequences in GenBank (http://blast.ncbi.nlm.nih.gov/Blast.cgi) using basic local alignment search tool (BLAST) analysis. The sequences of samples positive for *Borrelia crocidurae* by qPCR are shown in Figure 2. Phylogenetic trees were constructed using the maximum likelihood method with model selection determined by Molecular Evolutionary Genetics Analysis (MEGA) v7.0.26 software (Tamura K, 1993). The statistical support of the internal branches of the trees was assessed by bootstrapping with 1000 iterations.

### Ethical Considerations

RDTs were performed as part of routine care for patients with a fever at healthcare centres. This study did not involve any patients or collect any personal data. The samples were deidentified and collected at the healthcare centres. Regulatory approval for the use of malaria RDTs has been granted by the National Malaria Control Programme under number 064/MSDS-SG/NMCP.

## Results

### DNA was extracted from RDT cassettes (30 positive and 150 negative)

A total of 1496 malaria RDT samples (260 positive and 1236 negative) from febrile patients were tested for *Borrelia* spp. by real-time PCR. Of these, only (0.6%) 9/1496 RDTs were positive for both *ITS4* and *glpQ* qPCRs (**Figure 1**). *Borrelia crocidurae* DNA was detected in 5% of RDTs tested (9 out of 180 randomly selected), collected in the Kayes region. Querying all the sequences obtained against the GenBank database showed 99 to 100% coverage and 100% identity with the *B. crocidurae* sequence (JX119098) from Senegal. The phylogenetic tree shows that our sequences are close to those already found in Mali, Mauritania and Senegal (**Figure 2**).

**Figure 1.**
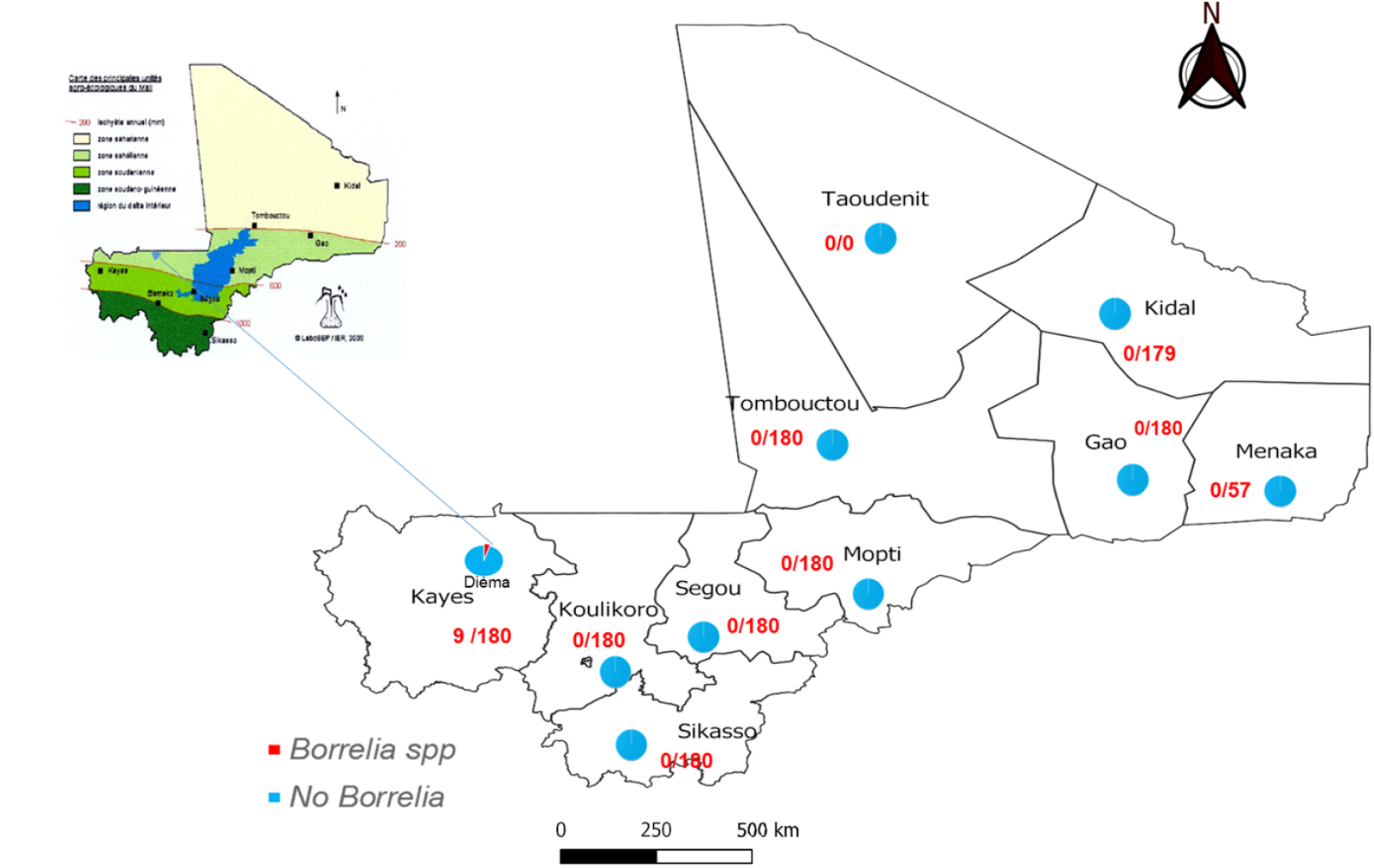
Map of Mali showing the number of malaria RDT cassettes tested in the denominator and the number positive in the numerator

**Figure 2.**
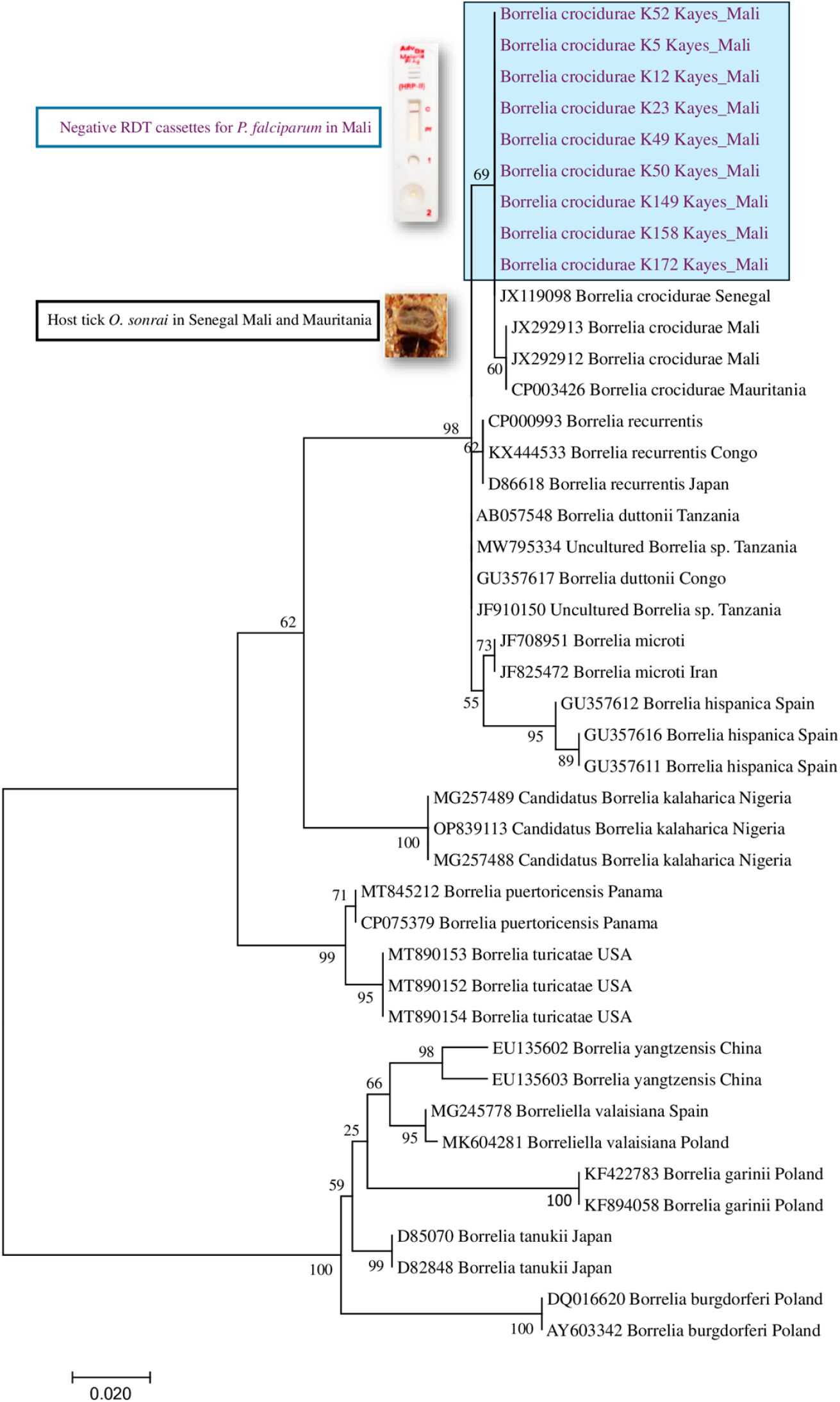
Phylogenetic tree (maximum likelihood method, bootstrap 1,000) of partial sequence data for the flagellin gene, (FlaB) of *B. crocidurae* identified from malaria RDTs.

Sequences of the *B. crocidurae FlaB* gene found in *O. sonrai* ticks and sequences from other *Borrelia* species were processed for comparison. The tree with the highest likelihood (-1092.19) is shown.

## Discussion

Tick-borne relapsing fever (TBRF) is the most common vector-borne bacterial human disease in West Africa. Between December 2007 and October 2011, a study conducted in 20 villages in southern Mali reported the presence of the vector *O. sonrai* in small mammals and ticks collected from rodent burrows and found that 17.3% of ticks were infected with spirochetes [17]. In this study, we found a relatively high prevalence of 6% *B. crocidurae* in negative malaria RDTs of febrile patients in a rural health centre in the Diéma health district, located in the Kayes region bordering Senegal and Mauritania, two *Borrelia*-endemic countries [2,37].

Most studies of TBRF in West Africa have been conducted in Senegal. Our results are comparable to those of Mediannikov et al. conducted in Senegal between June 2010 and October 2011, where 115 (7.3%) of 1,566 blood samples from febrile patients were positive for *B. crocidurae*. The highest proportion of positive samples was found in Niakhar with 19.1% (33/173) [38]. Patients aged 7 to 15 years were most affected: 13.5% (33/173). The proportion of *Borrelia*-positive samples was significantly higher in samples collected during the dry season (16.9%, 40/237) than during the rainy season (6.9%, 30/432). Several previous studies have demonstrated the influence of climate change on infectious diseases [21,39,40]. In December 2016, among febrile patients examined in one health centre and three health posts in Niakhar, Ngayokheme, Toucar and Diohine, respectively, cases of borreliosis accounted for 12% (94/800) of fever episodes, and all age groups were infected, with children and adolescents aged 8-14 and 22-28 years being most affected by the disease (16% and 18.4%). The increase in the number of cases demonstrates that TBRF is a real public health problem in Senegal and other neighbouring countries such as Mali. Contrary to the data from the previous study, the highest number of TBRF cases occurred in August [22], which also very often coincides with the period of high malaria transmission.

To avoid overdiagnosis of malaria cases, health care providers must follow the following guidelines for malaria case management [41], as the correct use of RDTs should optimise malaria diagnosis and prevent antimalarial drug resistance. RDTs are also considered a convenient source of DNA for molecular analysis [25]. In this context, a similar study by Ndiaye EHI et al. found a 7% prevalence of *B. crocidurae* in malaria RDTs collected in Senegal [7].

The *B. crocidurae* detected in our malaria-negative RDT DNA samples has previously been identified in *O. sonrai* ticks in Senegal, Mauritania and Mali [1,42]. Phylogenetic analysis of the flaB gene showed that the *Borrelia* in our study were similar to those previously found in Senegal [17,42]. The results presented in this study confirm the involvement of *B. crocidurae* causing TBRF in non-malaria fever cases in Mali.

## Conclusion

The surveillance system for vector-borne diseases such as borreliosis needs to be strengthened, and diagnostic tools for TBRF that are sensitive and much more specific for *B. crocidurae* need to be adopted to improve the management of this endemic zoonotic infection in Mali. Strengthening the skills of microscopists may be one way to improve the diagnosis of the disease.

## Author contributions

P.D., A.Z.D., M.A.T. and S.R. conceived the study and drafted the first version of the manuscrip**t**.

P.D., A.Z.D., P.M.G., L.B., and., A.D.A.J analyzed and interpreted the results.

A.Z.D., M.A.T. and S.R. contributed to the critical revision of the manuscript. All authors have read and approved the version

## Acknowledgements

This study was made possible by the academic and financial support of the Fondation Méditerranée Infection in Marseille, France, and the management of the National Malaria Control Program. We thank each and every one of you.

## Conflicts of interest

The authors declare no conflict of interest.

## Availability of data and materials

The authors confirm that the data supporting the results of this study are available in the manuscript and its supplementary information files.

## Notes

### Competing Interest Statement

The authors have declared no competing interest.

